# A collection of Predicted No-Effect Concentrations of human pharmaceuticals and their metabolites

**DOI:** 10.1101/2023.12.12.571257

**Authors:** Valentina Giunchi, Elisabetta Poluzzi

**Affiliations:** Department of Medical and Surgical Sciences, Alma Mater Studiorum – Università di Bologna, Bologna, Italy

## Abstract

The environmental impact of pharmaceuticals is a growing concern, necessitating methodologies for risk assessment. Current evaluation methods rely on comparing pharmaceutical concentrations in exposed environments with relevant animal or plant tolerance thresholds, often represented by Predicted No-Effect Concentration (PNEC) values. However, challenges arise from the limited accessibility and standardization of PNEC data. This study addresses these challenges by consolidating PNEC values from diverse sources into a unified and accessible tool. Investigated data sources were the NORMAN Ecotoxicology Database, the Swedish National Formulary of Drugs (FASS) website, the EU Watch List working documents, the European Public Assessment Reports (EPAR), the US EPA ECOTOX database, the UBA ETOX database, the WikiPharma database, the AstraZeneca documents, the European Chemical Agency registration dossiers, and the AMR Industry Alliance database. We retrieved 93,287 PNEC values associated with 92,850 substances, primarily medicines or their metabolites. Notably, 352 substances had more than one PNEC value, with the highest discrepancies often attributed to in-silico predicted values. The resulting database, available in the related OSF repository as a spreadsheet file, includes source information and processing scripts and is freely available for risk assessment analyses. While acknowledging limitations, future efforts should prioritize integrating additional data sources, addressing misspellings, and enhancing information on PNEC derivation. Collaboration in PNEC data collection is crucial for advancing collective knowledge in pharmaceutical risk assessment.

## Introduction

The assessment of the environmental impact of pharmaceuticals is gaining increasing attention within both regulatory and research frameworks. The most widely recognized method for evaluating this impact involves comparing the concentration of pharmaceuticals in a potentially exposed environment with the relevant tolerance thresholds of animals or plants. When it comes to pharmaceuticals intended for human use, surface water is the preferred environment for conducting environmental assessments. This preference stems from the fact that pharmaceuticals, once consumed, are excreted in urine and feces, subsequently finding their way into wastewater and, ultimately, surface water bodies.

While concentration measurements can be either sampled or estimated, the tolerance threshold usually consists of a Predicted No-Effect Concentration (PNEC). PNEC values are typically derived from in-vivo experiments. According to European legislation governing pharmaceutical marketing approval, these experiments should be conducted following the internationally standardized guidelines provided by the Organization for Economic Co-operation and Development (OECD)^1^. However, it’s important to note that OECD standard tests were originally developed for assessing industrial chemicals, and they may not be entirely suitable for pharmaceutical testing^2,3^. Recognizing these limitations, researchers have already proposed changes to European Medicines Agency (EMA) legislation, suggesting the utilization of existing ecotoxicity data instead of mandating new testing. This transition would yield several benefits, including more efficient resource utilization by avoiding unnecessary economic and scientific efforts, as well as aligning with ethical considerations by minimizing redundant animal testing^3^. In addition to the marketing authorization stage, PNEC values hold significance in the post-marketing setting. Indeed, when confronted with measured or predicted water concentrations of pharmaceuticals, PNEC values play a crucial role in assessing whether these concentrations could potentially pose an environmental risk. Recent studies have highlighted the challenge of obtaining PNEC values, thereby rendering it impossible to derive a risk assessment for numerous pharmaceuticals. This issue is compounded by the fact that even when PNEC values are accessible, their retrieval can often be time-consuming. For instance, in the case of the European Public Assessment Reports (EPAR), information must be extracted from PDFs organized by brand name. Similarly, the Swedish National Formulary of Drugs (FASS) website arranges information on webpages classified by brand name and presented in the Swedish language, further adding to the complexity^4,5^.

The objective of this study is to review existing and publicly accessible sources of PNEC values and consolidate them into a single, easily accessible tool. This initiative aims to enhance the utilization of already available results of ecotoxicological tests, benefiting both industrial applications (such as the pre-marketing authorization stage, if permitted) and research, as well as post-marketing regulatory contexts. The approach undertaken seeks complete transparency in its methodology, with the intention of fostering meaningful dialogues among diverse experts regarding the inclusion of additional sources and the criteria for selecting the most appropriate PNEC values in cases where disparate studies yield differing results. The authors’ intention is not to supplant the existing sources of information, which are essential and represent the foundation of PNEC knowledge thanks to the efforts of their creators. This endeavor solely seeks to offer a more inclusive and user-friendly approach to accessing these data.

## Methods

Sources of PNEC data were identified through web research and by referencing articles. Unfortunately, alternative methods such as searching through Web of Science or PubMed were found to be less suitable due to the frequent occurrence of PNEC sources being institutional webpages or databases hosted on research websites.

In the following, each data source utilized to construct the collection of PNEC values is delineated, along with an account of the standardization steps undertaken.

### NORMAN Ecotoxicology Database – Lowest PNECs

The NORMAN Ecotoxicology Database is maintained by the NORMAN network, a consortium of research centers and laboratories established in 2005 with financial support from the European Commission. Its primary objective is to monitor emerging environmental substances. The Lowest PNECs section of the NORMAN Ecotoxicology Database (accessible at www.norman-network.com/nds/ecotox/lowestPnecsIndex.php) compiles PNEC values related to long-term exposure. These values are gathered from publicly available sources, such as published literature and databases. Additionally, in-silico methods, such as quantitative structure-activity relationships (QSAR), are employed to derive these values as well^6^. It’s important to note that the reliability of in-silico-derived PNEC values might not always be as high as those obtained through in-vivo methods^5^. Nevertheless, this study includes both types of values and indicates whether a value is predicted in-silico. This approach allows researchers to conduct risk assessments even in the absence of other available PNEC values and empowers them to evaluate the appropriateness of using in-silico predictions for their assessments.

The NORMAN Lowest PNECs database was downloaded as a CSV file on July 22nd, 2023, and subsequently pre-processed in R to create a database that could be effectively merged with the other data sources. The steps involved in this pre-processing are illustrated in **Figure 1**.

**Figure 1.**
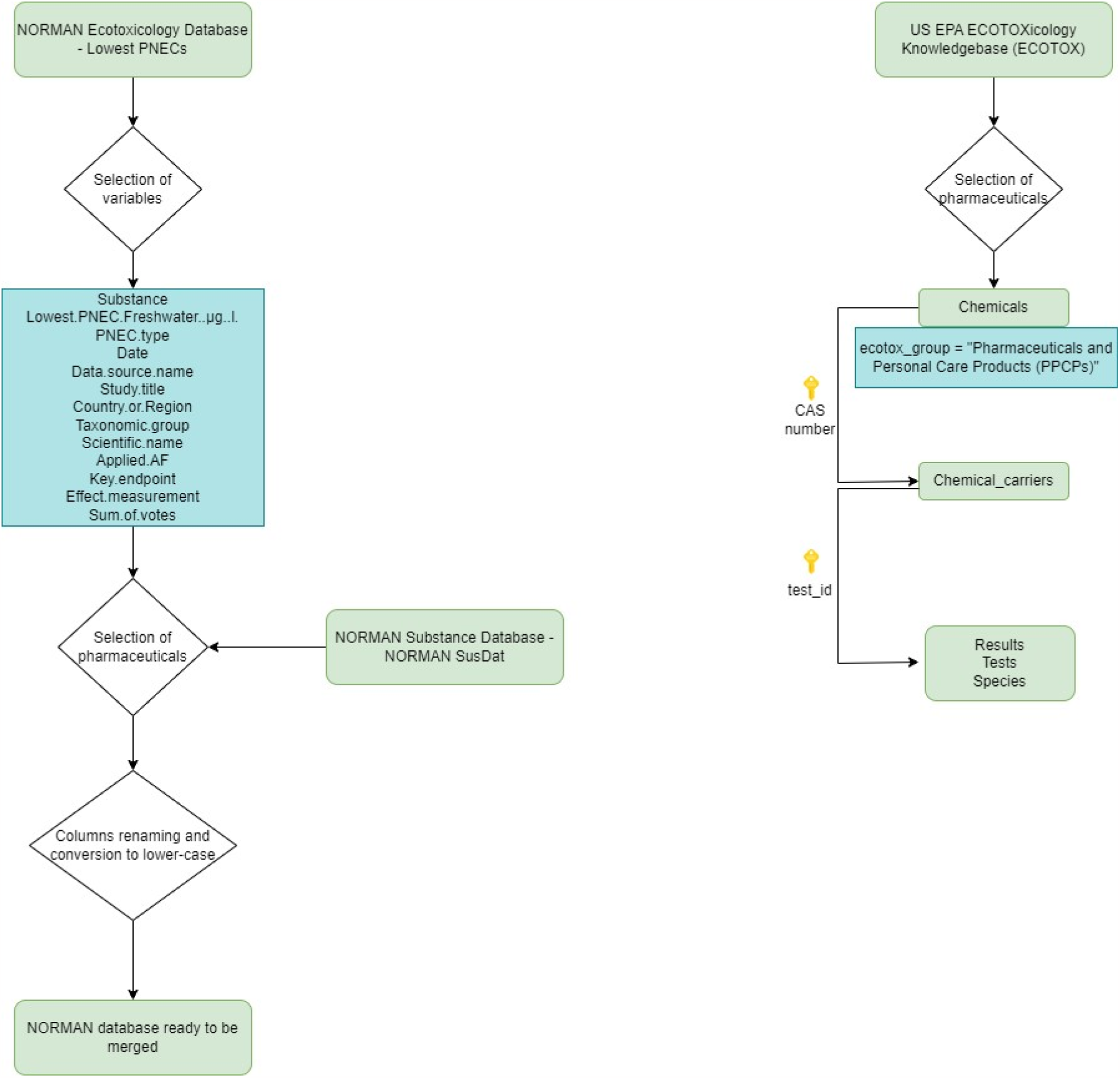
Pipeline of steps followed during NORMAN Lowest PNECs pre-processing on the left and US-EPA ECOTOX on the right.

### Swedish National Formulary of Drugs (FASS) website

The Swedish National Formulary of Drugs (FASS) website (accessible at www.fass.se/LIF/startpage) hosts environmental risk assessment documents submitted by manufacturers during the marketing authorization stages for medicinal products sold in Sweden. Occasionally, information is also available for imported and withdrawn products. Notably, there exists no comprehensive list of medicinal products for which environmental information is accessible, and this information is stored according to brand names. To gather this data, each pharmaceutical name (defined by the Active Pharmaceutical Ingredient - API) with an ATC code was searched, and all corresponding brand name webpages were explored to identify the presence of environmental information. Data extraction was undertaken by a single researcher from April to June 2023, and its accuracy was subsequently verified by two other researchers between July 22nd and August 4th, 2023.

Additionally, the Norwegian Pharmaceutical Product Compendium (Felleskatalogen) also furnishes details regarding the environmental impact of pharmaceuticals. However, for the purpose of this analysis, it was not considered due to its assertion that it employs the same ecotoxicological information contained in FASS^7^.

### European Watch List working documents

The European Commission established the Directive 2008/105/EC, which introduced a list of chemical substances to be monitored in Member States’ surface waters due to potential environmental risks^8^. The initial list of substances was defined in 2015, and since then, it has undergone four subsequent updates. In general, a substance becomes eligible for inclusion if there is ecotoxicological evidence suggesting it could pose an environmental risk, but the scientific literature alone is insufficient to ascertain whether its actual presence in surface waters constitutes a risk. Once sufficient information is gathered, a substance can be removed from the list. In the four updates conducted thus far, some pharmaceuticals have been added to the list.

To access PNEC values for included and candidate pharmaceuticals that were used during the substance selection process, we thoroughly reviewed the Watch List working documents^9–12^. This data extraction was undertaken by one researcher and subsequently verified by another in April 2023.

### European Public Assessment Report (EPAR) documents

In 2006, the EMA introduced the Environmental Risk Assessment (ERA) as a component of human medicines EPAR for manufacturers undergoing the marketing authorization procedure. Specifically, ERA guidelines, which underwent revision in 2018, mandate a Phase I assessment based solely on the expected utilization of the pharmaceutical in the market, along with a Phase II assessment focused on risk characterization. Consequently, for pharmaceuticals requiring Phase II assessment, manufacturers are obligated to conduct in-vivo tests following OECD guidelines and report the results in the ERA^1^. Despite the longevity of this obligation since 2006 and its reinforcement in 2018, the number of EPAR featuring complete and transparent ERA remains limited^13^.

Extracting information from the EPAR documents is challenged by the fact that they are arranged based on their last modification date, leading to continuous updates. This updating not only includes new products but also minor modifications to existing ones. Furthermore, due to the primary focus of EPAR on providing environmental information for new products, the reliance on the ATC classification list may not always be effective, as some products might not yet have an assigned ATC code. Additionally, there are legacy medicines approved before 2006, for which it is already established that no environmental information exists in EPAR. To retrieve ecotoxicity information from EPAR, two researchers conducted searches on April 27th and 30th, 2023, as well as on May 6th, 9th, 11th, 13th, 15th, 18th, 19th, and 23rd, 2023, focusing on the newest or most recently updated products and the results have been checked by another researcher on August 21st, 2023. The complete list for which ERA was searched is reported in the OSF repository together with extraction information.

### US EPA ECOTOX

The ECOTOXicology Knowledgebase (ECOTOX) is a relational database maintained by the United States Environmental Protection Agency (US EPA). This comprehensive database houses data on chemical toxicity for aquatic organisms, terrestrial plants, and wildlife^14^. The data were downloaded as ASCII files on May 14th, 2023, and subsequently processed using R (**Figure 1**).

### UBA ETOX

The German Environment Agency (Umweltbundesamt – UBA) oversees ETOX, the Information System for Ecotoxicology and Environmental Quality Targets. This system offers comprehensive information on ecotoxicology tests conducted in aquatic and terrestrial environments, with a focus on substances pertinent to the evaluation of surface water pollution. Its primary aim is to facilitate the establishment of environmental quality standards in alignment with the Water Framework Directive 60/2000/EC^15^.

In ETOX, a comprehensive download of the complete dataset is not available; instead, users can conduct searches using a graphical user interface. Specifically, there are two distinct databases available for searching: the “Quality Target” database, which contains quality measures like PNEC, and the “Effect Data” database, which houses ecotoxicological endpoints such as NOEC or EC50. In regard to the first dataset, we conducted a data selection process by specifying “Medium/Matrix” as “WATER” and “Designation” as “PNEC” on August 16th, 2023. This approach limited the number of extracted values and allowed for their export in CSV format. Subsequently, one researcher carried out a manual selection of the substances to be included, specifically pharmaceuticals. The search of the second database proved to be more challenging due to our potential interest in a variety of effects and endpoints and was hindered by technical issues.

### AstraZeneca data

In 2017, AstraZeneca’s pharmaceutical manufacturing division released a document that presented the ERA of their products. These assessments were compiled from submissions made to the EMA or other regulatory agencies, or, in the case of legacy medicines, were sourced from available published literature^16^. We extracted the PNEC values reported here by accessing pharmaceutical-specific PDFs.

### ECHA Registration Dossiers

The European Chemicals Agency (ECHA) maintains a web-based database containing information on all registered substances. This database is particularly comprehensive, encompassing environmental data including details on aquatic, terrestrial, and sediment toxicity, as well as information about biodegradation, bioaccumulation, transport, and distribution^17^. We conducted a search for pharmaceuticals on August 22nd, 2023 using the search menu, specifically selecting “uses and exposure”, followed by “product category”, and then “PC29 pharmaceuticals”. However, it’s important to note that the information retrieved doesn’t encompass pharmaceuticals exclusively, it also includes medicines’ metabolites, additives, and excipients.

### WikiPharma

WikiPharma is a database containing regularly updated ecotoxicity data for pharmaceuticals. It was developed as part of the Swedish research program MistraPharma, which received funding from the Swedish Foundation for Strategic Environmental Research (Mistra). The data included were curated by searching for ecotoxicity data in scientific databases such as CSA, ScienceDirect, and PubMed, as well as utilizing the Google Scholar search engine and relevant books. The provided information includes details about the pharmaceutical active ingredients, the test method employed, the tested species, the sex, and the age or life stage of the organism, the number of organisms involved, API doses, exposure time, study period, route of exposure, test results, and critical effects (endpoints)^18^. While WikiPharma is intended to be accessible at www.wikipharma.org, it is important to note that as of the current date (November 16th, 2023), the website is unavailable.

### AMR Industry Alliance – Responsible Manufacturing

The Antimicrobial Resistance (AMR) Industry Alliance published a set of PNEC values for antibiotics, driven by the recognition that the environmental toxicity of some of these substances remained incompletely explored. They introduced both an “environmental PNEC” (PNEC-ENV) and an “inhibitory PNEC” (PNEC-MIC). The latter refers to the lowest concentration of an antibiotic that completely inhibits visible growth of a specific bacterial strain after 24 hours of incubation^19,20^. For this study, we extracted these data on August 14th, 2023, focusing exclusively on the “PNEC-ENV”. It’s important to note that we choose these PNEC values as they were derived from toxicity endpoint data, with an assessment factor applied, consistent with European guidance.

## Results

A total of 93,287 PNEC values associated with 92,850 substances were retrieved. The search primarily targeted medicines and their metabolites. However, despite efforts to restrict queries to these substances whenever possible, some additional chemical compounds were included. To account for the heterogeneity in the terminology used to identify substances in the data sources included, a new field named “Substance_transl” was introduced alongside the original substance names recorded in the “Substance” field. Currently, the “Substance_transl” field is empty, offering researchers the opportunity to translate substances as needed without losing the original information. This flexibility allows for different choices based on research-specific requirements. The presented results were obtained through a semi-automatic standardization process, involving the correction of typographical errors and misspellings (e.g., *aalizapride* to *alizapride*), the conversion of Greek alphabet letters to their corresponding Latin spellings, and the translation of non-English terms into English (e.g., *acenocumarolo* to *acenocoumarol*). Other fields included in the retrieved database are the PNEC (expressed in μg/L) and the PNEC type, which applies exclusively to data derived from NORMAN. As mentioned earlier, NORMAN comprises both in-vivo and in-silico-derived PNEC, with the latter referred to as predicted PNEC. Additionally, the database includes information such as the date, country, species (group, scientific name, and taxonomic group) on which the test was performed, the endpoint, the assessment factor, and the effect considered for PNEC derivation. Furthermore, details about the source from which the PNEC originates, and information related to the NORMAN database, such as the monitoring program and the credibility of the source, are also included

### Multiple PNEC values

Among the retrieved substances, 352 (0.38%) have more than one associated PNEC value. The substances with the highest number of PNEC values retrieved are trimethoprim (N=8), sulfamethoxazole (N=8), metformin (N=6), and ciprofloxacin (N=6).

There are 298 pharmaceuticals (0.32% of the total) that possess two PNEC values. Most of the discrepancies between these paired values are attributed to one of them being in-silico predicted, specifically derived using the NORMAN QSAR approach (**Figure 2**). Moreover, 56 pharmaceuticals show no difference between their paired PNEC values, signifying duplication. Excluding the in-silico predicted values, which are generally considered less reliable than in-vivo determinations, the highest differences were observed for iopromide (10,000 μg/L vs. 20,000 μg/L), terbutaline (3,000 μg/L vs. 240 μg/L), and topiramate (930 μg/L vs. 93 μg/L).

**Figure 2.**
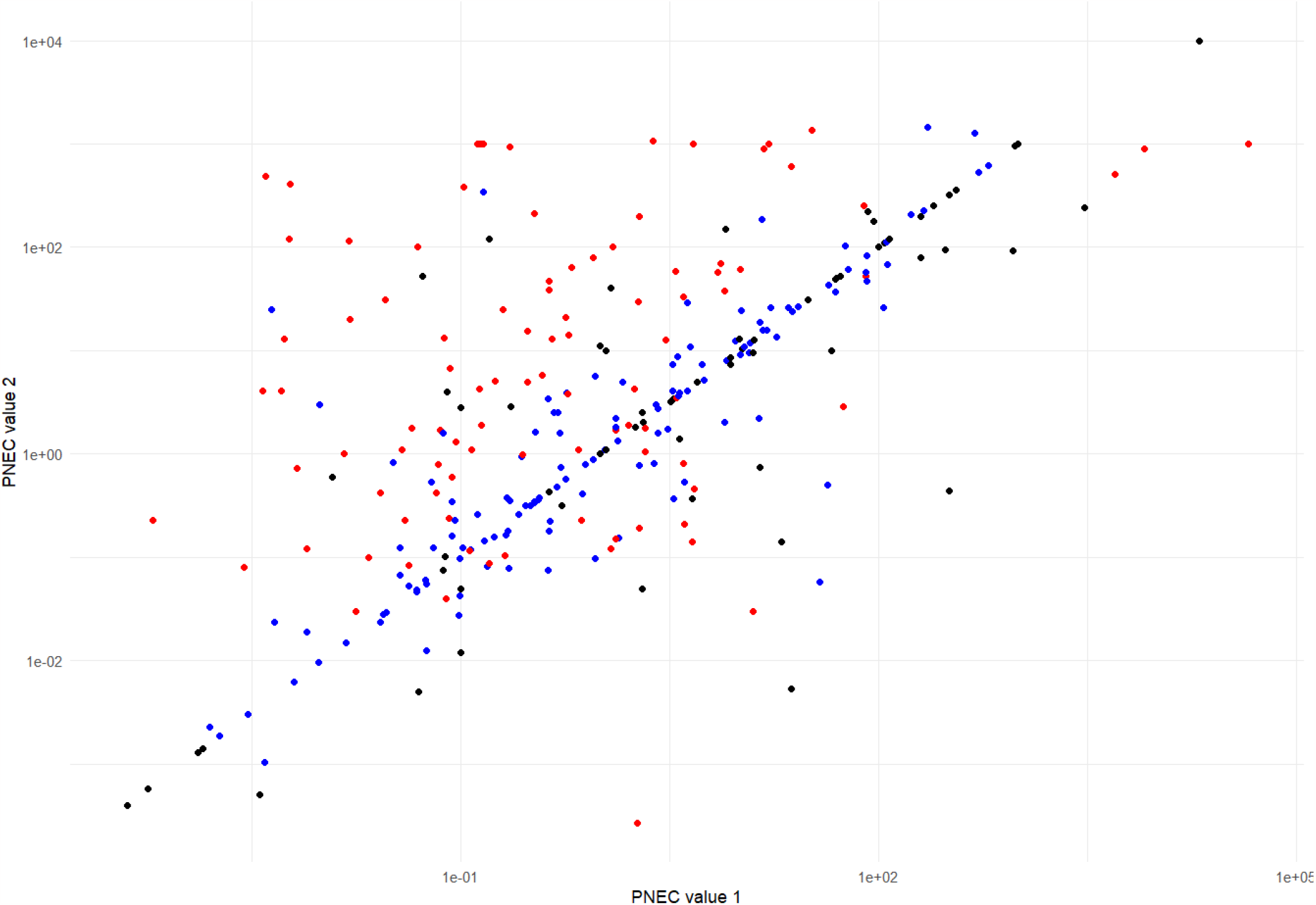
PNEC values for pharmaceuticals with two retrieved values are visualized with two axes representing each PNEC value. Dots are displayed in black if both PNEC values are in-vivo derived, in blue if they are both in-silico derived, and in red if one PNEC is derived in-vivo and the other in-silico.

There are 54 pharmaceuticals (0.06% of the total) with more than two PNEC values. Among them, seven pharmaceuticals had completely overlapping values, resulting in a standard deviation (SD) of zero. Specifically, these substances —dapagliflozin, esomeprazole, ibrutinib, quetiapine, rosuvastatin, ticagrelor, and zolmitriptan— yielded identical values across different sources, even though minor discrepancies in effects data might exist (which could be attributed to processing variations). Conversely, six pharmaceuticals exhibited multiple values with notable variability (SD > 100 μg/L). These were gabapentin (SD = 571.57, N = 3), tenofovir alafenamide (SD = 550.74, N = 3), ramipril (SD = 519.62, N = 3), metformin (SD = 493.38, N = 6), lidocaine (SD = 317.47, N = 3), and trimethoprim (SD = 102.29, N = 8) (**Figure 3**). It’s noteworthy that, in the case of tenofovir alafenamide, one of the PNEC values with the highest discordance is an in-silico predicted value sourced from the NORMAN database (**Figure 4**).

**Figure 3.**
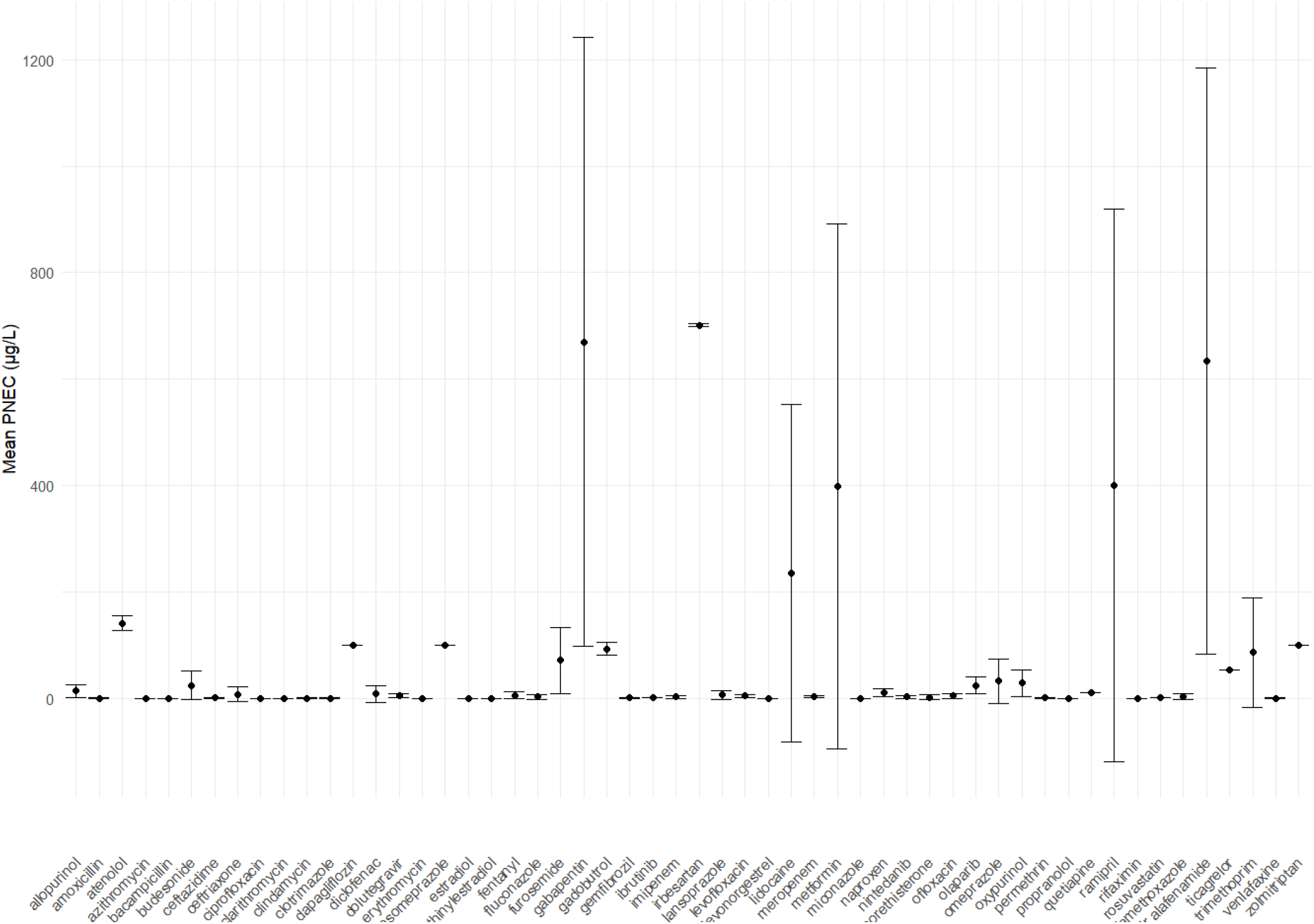
Means and standard deviations of PNEC of pharmaceuticals with more than two PNEC values. Dots represent the mean value of retrieved PNECs, bars represent the mean value minus and plus the standard deviation of each pharmaceutical.

**Figure 4.**
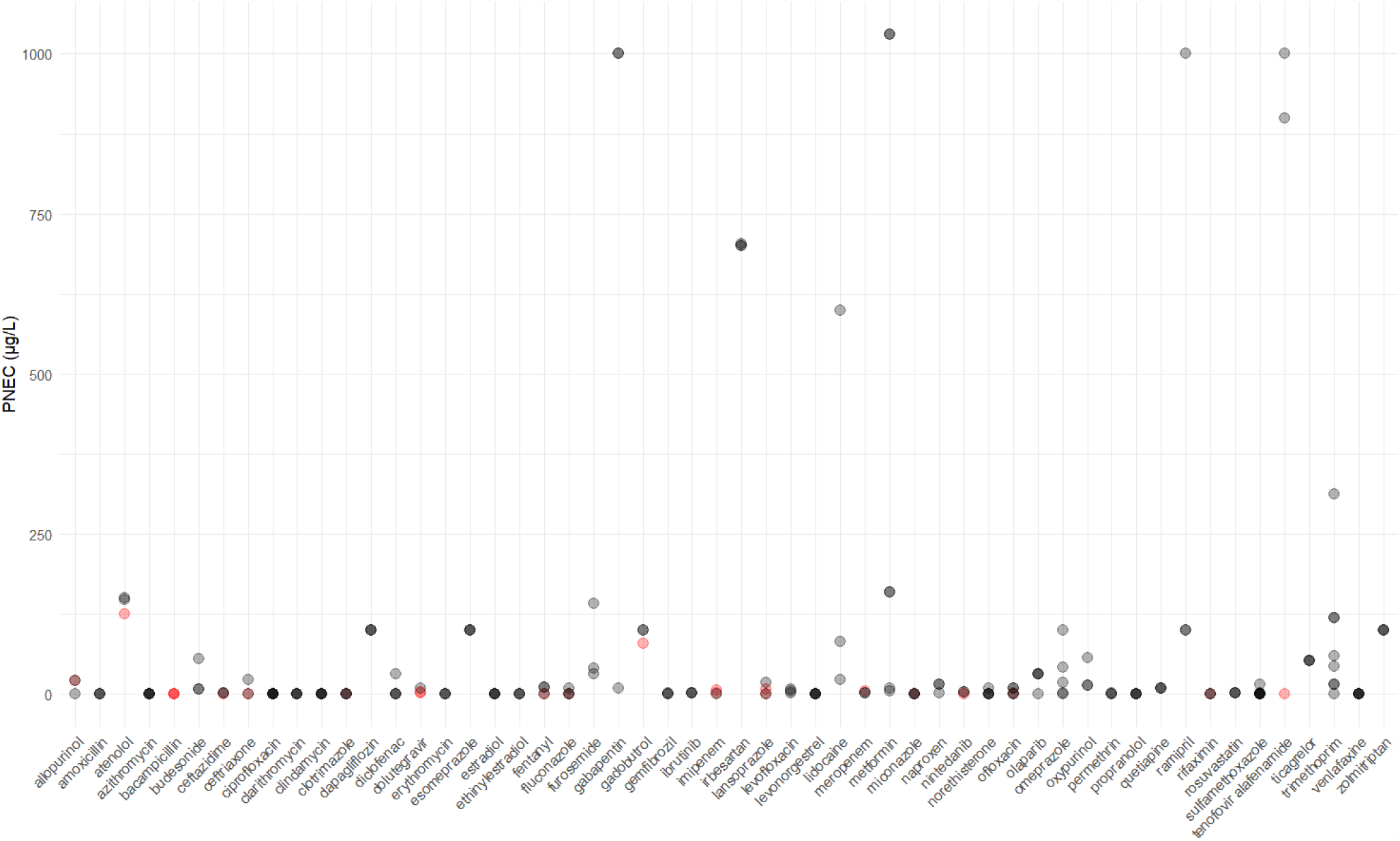
Pharmaceuticals with more than two PNEC values are graphically represented. The x-axis denotes the pharmaceutical, and the y-axis shows corresponding PNEC values. In-vivo-derived PNEC values (freshwater) are depicted in gray, while those derived through in-silico methods (e.g., QSAR) are in red. Darker shading indicates multiple occurrences of the same PNEC value in the dataset.

### Obstacles in data collection

Among the consulted PNEC data sources, it was not possible to access the MistraPharma database, since the related website is currently inactive. Additionally, challenges arose when attempting to extract data from EPAR documents, the ECHA Registration Dossiers, and the UBA ETOX “Effect Data” database. For EPAR, the number of documents consulted is limited, with the intention of expanding consultation efforts in the future. Similarly, further efforts are needed for the review of the ECHA and the UBA data sources. The extraction of data from these two databases was hindered by the heterogeneity of the data, and setting up a precise research query proved to be challenging.

## Discussion

The aim of this work was to map the existing sources of PNEC data and gather them to create a useful resource for risk analyses. This database, shared as a spreadsheet file along with the source files and the R script used for its creation, is publicly accessible at the project OSF repository (DOI: 10.17605/OSF.IO/XTG8Z).

This work acknowledges some limitations and identifies future challenges. Certain data sources chosen for PNEC extraction, such as WikiPharma, were not accessible, while others like EPAR documents, ECHA, and UBA “Effect Data” databases were not easily accessible for information extraction. Future efforts should focus on integrating these data sources with the already collected data. Additionally, an explorative analysis of the created database revealed the presence of some misspellings or inconsistencies in substance names that need to be addressed. Furthermore, future research in PNEC collection should consider including more information about PNEC derivation, such as whether the test was performed following specific directives (e.g., OECD guidelines). At present, when multiple PNEC values are available, and it is not possible to distinguish one from others is possible based on the reliability of their derivation, environmental risk assessment should be based on the lowest PNEC value to define the worst-case scenario. Additional efforts should be directed towards broadening data sources, beginning with a comprehensive literature review, to integrate into our database. This approach may uncover valuable single studies that provide PNEC values, particularly for new or understudied pharmaceuticals.

Collaboration on PNEC data is crucial to furnish researchers with a readily accessible database. The importance of pharmaceutical risk assessment in surface water is growing, both from regulatory and academic perspectives. Consequently, it becomes imperative to harmonize efforts and foster collaboration to build and share collective knowledge.

## Data availability statement

Data used, R script with analysis code and the final dataset are publicly available in the following OSF repository: https://osf.io/xtg8z/ (DOI: 10.17605/OSF.IO/XTG8Z)

## Acknowledgements

V.G. was supported by EU funds (Programma Operativo Nazionale Italian funds for green and innovative research based on European Structural and investment Funds). E.P. was supported by institutional research funds (Ricerca Fondamentale Orientata).

We would like to express our gratitude to Federica Fiore and Giorgia Maria Lari (Biological Sciences BSc trainees during 2023 spring semester) for their assistance in extracting and revising PNEC values, and to our colleague Michele Fusaroli for revising the R code.

## Author contributions

The study was conceptualized by both authors. V.G. collected and manipulated the data to create the final dataset, and subsequently wrote the original draft. E.P. edited and revised the manuscript. Both authors reviewed and approved the final version.

## Conflict of interest statement

The authors declare that they have no conflicts of interest.

## Notes

https://osf.io/xtg8z/

### Competing Interest Statement

The authors have declared no competing interest.

## References

1. EMA. Revised guideline to assess risk of human medicines for the environment. European Medicines Agency. Published November 30, 2018. Accessed August 5, 2023. https://www.ema.europa.eu/en/news/revised-guideline-assess-risk-human-medicines-environment

2. Bound JP, Voulvoulis N. Pharmaceuticals in the aquatic environment––a comparison of risk assessment strategies. Chemosphere. 2004;56(11):1143–1155. doi:10.1016/j.chemosphere.2004.05.010

3. Ågerstrand M, Berg C, Björlenius B, et al. Improving Environmental Risk Assessment of Human Pharmaceuticals. Environ Sci Technol. 2015;49(9):5336–5345. doi:10.1021/acs.est.5b00302

4. Giunchi V, Fusaroli M, Linder E, et al. The environmental impact of pharmaceuticals in Italy: Integrating healthcare and eco-toxicological data to assess and potentially mitigate their diffusion to water supplies. Br J Clin Pharmacol. 2023;89(7):2020–2027. doi:10.1111/bcp.15761

5. Welch SA, Moe SJ, Sharikabad MN, Tollefsen KE, Olsen K, Grung M. Predicting Environmental Risks of Pharmaceuticals from Wholesale Data: An Example from Norway. Environ Toxicol Chem. n/a(n/a). doi:10.1002/etc.5702

6. NORMAN Ecotoxicology Database. Accessed August 5, 2023. https://www.norman-network.com/nds/ecotox/

7. Felleskatalogen. Legemidler og miljø. Accessed August 20, 2023. https://www.felleskatalogen.no/medisin/miljo/innledning

8. Commission Implementing Decision (EU) 2015/495 establishing a watch list of substances for Union-wide monitoring in the field of water policy pursuant to Directive 2008/105/EC of the European Parliament and of the Council. Accessed August 20, 2023. https://www.ecolex.org/details/legislation/commission-implementing-decision-eu-2015495-establishing-a-watch-list-of-substances-for-union-wide-monitoring-in-the-field-of-water-policy-pursuant-to-directive-2008105ec-of-the-european-parliament-and-of-the-council-lex-faoc142647/

9. Negrão DCR, Ceriani L, Ippolito A, Lettieri T. Development of the First Watch List under the Environmental Quality Standards Directive. JRC Publications Repository. doi:10.2788/101376

10. Loos R, Marinov D, Sanseverino I, Napierska D, Lettieri T. Review of the 1st Watch List under the Water Framework Directive and recommendations for the 2nd Watch List. JRC Publications Repository. doi:10.2760/614367

11. Selection of substances for the 3rd Watch List under the Water Framework Directive - Publications Office of the EU. Accessed August 20, 2023. https://op.europa.eu/en/publication-detail/-/publication/a2ab9f86-d140-11ea-adf7-01aa75ed71a1/language-en

12. Gomez CL, Marinov D, Sanseverino I, et al. Selection of substances for the 4th Watch List under the Water Framework Directive. JRC Publications Repository. doi:10.2760/01939

13. Health Care Without Harm. Recommendations for greener human medicines in the revision of the EU general pharmaceuticals legislation. Published online March 2022.

14. Olker JH, Elonen CM, Pilli A, et al. The ECOTOXicology Knowledgebase: A Curated Database of Ecologically Relevant Toxicity Tests to Support Environmental Research and Risk Assessment. Environ Toxicol Chem. 2022;41(6):1520–1539. doi:10.1002/etc.5324

15. ETOX: Information System Ecotoxicology and Environmental Quality Targets. Accessed August 21, 2023. https://webetox.uba.de/webETOX/index.do?language=en&path=index

16. AstraZeneca. AstraZeneca’s Environmental Risk Summaries. Published online 2017. Accessed August 22, 2023. https://www.astrazeneca.com/content/dam/az/PDF/2017/Environmental_risk_data_relating_to_our_medicines.pdf

17. Registered substances - ECHA. Accessed August 22, 2023. https://echa.europa.eu/en/information-on-chemicals/registered-substances

18. Molander L, Ågerstrand M, Rudén C. WikiPharma – A freely available, easily accessible, interactive and comprehensive database for environmental effect data for pharmaceuticals. Regul Toxicol Pharmacol. Published online September 1, 2009:367-371. doi:10.1016/j.yrtph.2009.08.009

19. Responsible Manufacturing. AMR Industry Alliance. Accessed August 22, 2023. https://www.amrindustryalliance.org/shared-goals/common-antibiotic-manufacturing-framework/

20. Vestel J, Caldwell DJ, Tell J, et al. Default predicted no-effect target concentrations for antibiotics in the absence of data for the protection against antibiotic resistance and environmental toxicity. Integr Environ Assess Manag. 2022;18(4):863–867. doi:10.1002/ieam.4560

